# Analytic representation of inhomogeneous-resolution maps of three-dimensional scalar fields

**DOI:** 10.1101/2022.03.28.486044

**Authors:** Alexandre G. Urzhumtsev, Vladimir Y. Lunin

## Abstract

Refinement of macromolecular atomic models versus experimental maps in cryo-electron microscopy and crystallography is a critical step in structure solution. For an appropriate comparison, model maps should mimic imperfections of the experimental ones, mainly atomic disorder and its limited resolution, often inhomogeneous over the molecule. We construct these model maps as a sum of atomic contributions expressed through a specially designed function describing a solitary spherical wave. Thanks to this function, atomic contributions analytically depend on both an atomic disorder and the local resolution, a value associated now with each atom. Such fully analytic dependence of inhomogeneous-resolution map values on model parameters permits an efficient refinement of all these parameters together and, beyond structural biology, opens a way to solve similar problems in other research domains.

**One-Sentence Summary:** An analytic decomposition of 3D-oscillating functions results in efficient tools to calculate maps and refine atomic models.

---

In this work, we propose a universal computational tool for a variety of projects that refer to maps of a limited resolution which may vary over space. For example, macromolecular atomic models (*1*) are obtained using experimental, or experimentally-based maps of spatial scalar fields such as an electrostatic scattering potential in cryo-electron microscopy (cryo-EM) or electron or nuclear density distributions in macromolecular crystallography (MX). Thanking to the “resolution revolution” in cryo-EM (*2*) and to a recent progress in structure prediction of protein components of macromolecular complexes (*3*), these maps become key tools to correct and refine initial atomic models suggested by these methods and to validate the results (*4*-*7*). Due to experimental features, the maps suffer from dynamic and static disorder and are available at some limited resolution which often vary from one macromolecular region to another (*8*) (Fig. 1). In order to refine an available model, one defines a respective score function comparing a map calculated from that model with the experimental one. For an appropriate quantitative comparison, the model map should mimic the imperfections of the experimental map. If the model map values are expressed analytically through the model parameters, this drastically simplifies the map calculation and the model optimization in general. We suggest a method to calculate such model maps in every point extending the concept of a local resolution (*8,9*) further and presenting the image of every atom *n* in the map with its own resolution *D*_*n*_. We have found an explicit expression for the map values by a developed analytic function of the atomic position **r**_*n*_, of its displacement parameter *B*_*n*_ and of individual resolution *D*_*n*_. This expression allows an accurate calculation of the inhomogeneous resolution map in a single run. Moreover, it allows simple analytic expressions for the gradient of the score function that rules the refinement of the model parameters increasing its efficiency. Finally, the *D*_*n*_ values can be refined together with other parameters, be reported with other atomic parameters such as its coordinates and *B*_*n*_ and deposited, in case of macromolecular models, in data bases such as PDB (*1*).

**Fig. 1.**
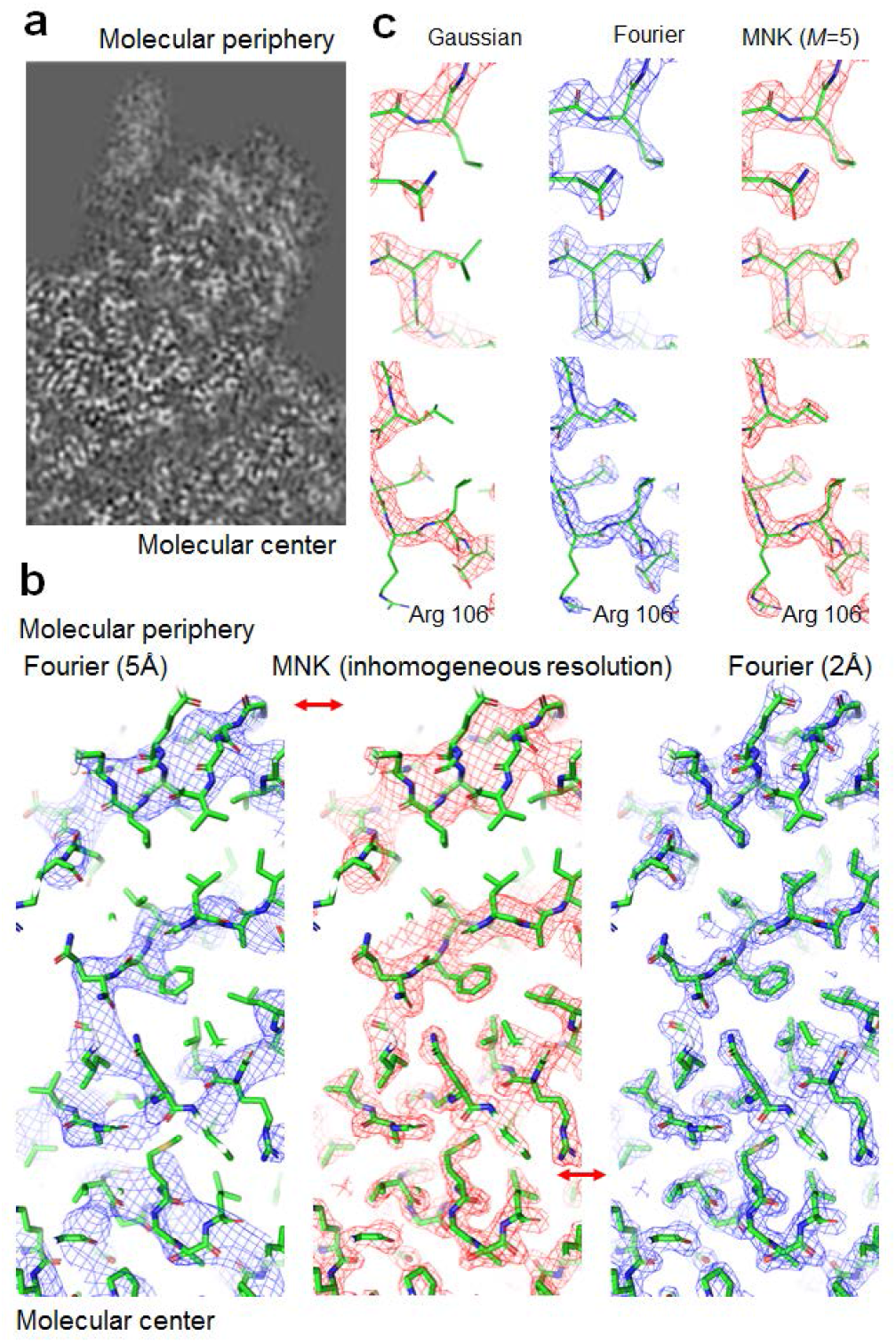
Maps of inhomogeneous resolution. (**a**) A fragment of a cryo-EM map calculated with a ‘home’ model (*26*) (EMDB 4261) illustrates a decrease of the local resolution from the molecular center to the periphery. (**b**) A *Ω*-map of an inhomogeneous resolution varying from 2 Å in the molecular center to 5 Å at the molecular periphery calculated in a single run by the shell-decomposition (middle), and the maps calculated by the Fourier procedure with the resolution of 5 Å (left) and 2 Å (right). Arrows mark the similarity of different parts of the *Ω*-map with the Fourier maps of different resolution. (**c**) Fragments of the 2 Å resolution maps contoured to show an equal volume (*27*). The map in the middle was calculated by the standard Fourier procedure. The left-hand map calculated as the sum of the Gaussian approximation to the atomic images (no ripples included) poorly reveals the density for some side chains. The right-hand map was calculated by the shell-decomposition (3) with *M* = 5 and reproduces the Fourier map correctly. A poor density for Arg106 residue in the *Ω*- and Fourier maps is due to large values of the displacement parameter of its atoms. The figure was prepared with *PyMol* (*28*).

Since mathematically the problem and its solution are the same for all relevant scalar fields, in what follows we will use the terms ‘density’ for the field, *ρ*(**r**), and an ‘atomic density’ for a contribution 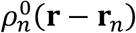 of a particular atom to this field. These atomic densities are usually described by spherically symmetric analytic functions 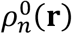 rapidly decreasing with distance, such as Gaussians or their combinations (*10,11*). In practice, they are cut at a relatively short distance, 2.5 – 3 Å. The maps *ρ*^*d*^(**r**) of these densities, to be compared with an experimental one, can be also seen as sums of the atomic contributions, 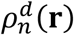, which this time we call atomic images in a given map and which should reproduce map distortions due to both their principal sources. While the atomic disorder blurs atomic densities, the resolution cut-off, beside this, generates Fourier ripples in the resulted image (Fig. 2a,b). Thus, the map distortions due to these two sources are not really equivalent (*12*). The ripples are seen as spherical waves of a slowly decreasing amplitude; they significantly contribute to the map quite far away from the atomic center (Fig. 1c).

**Fig. 2.**
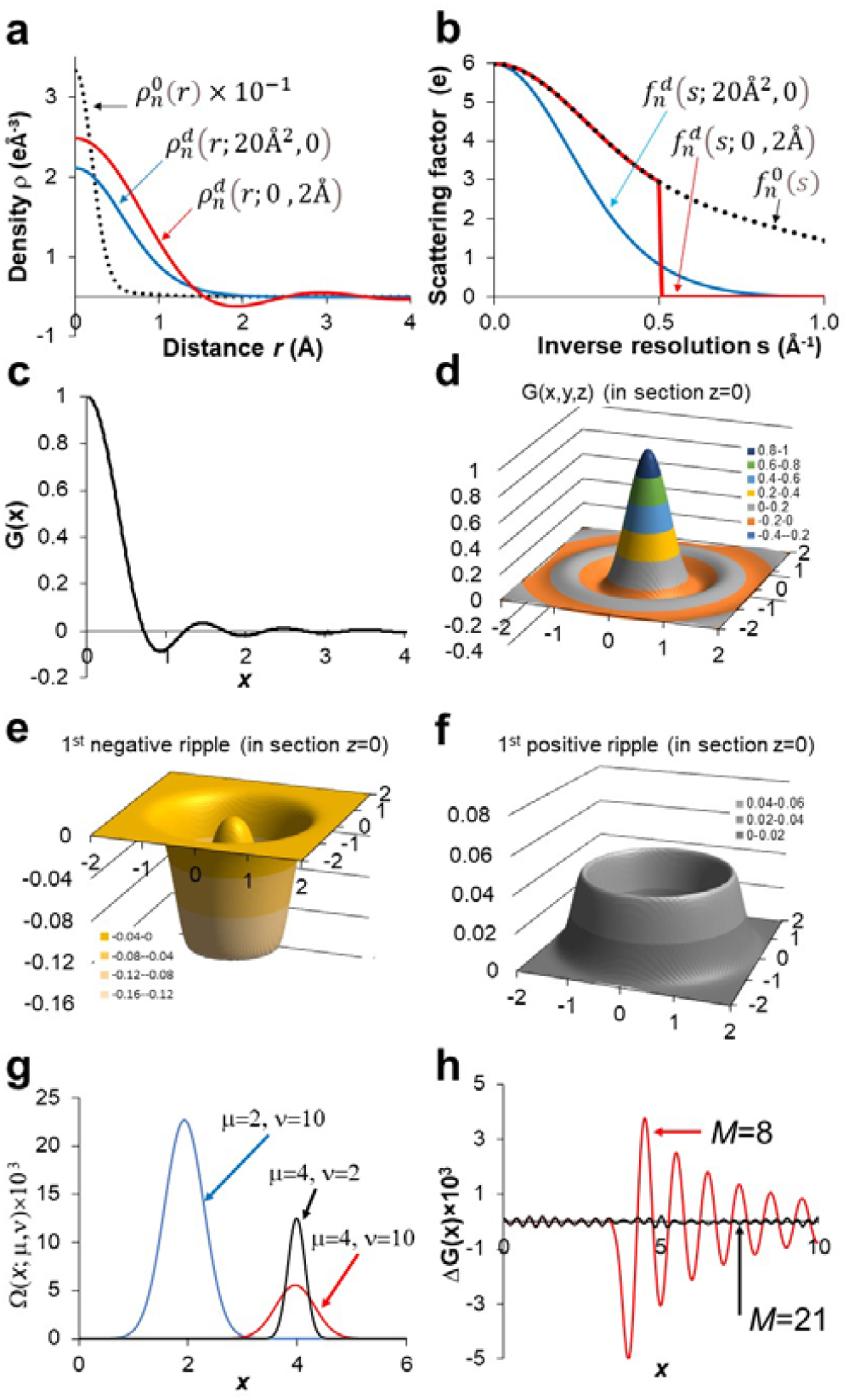
Atomic images and their decomposition. (**a**) The electron density distributions for an immobile carbon atom and its images affected by a disorder and a resolution cutoff. (**b**) Corresponding scattering functions (Fourier transform of the densities shown in Fig.2a). (**c**) The radial part of the interference function *G*(**x**) and (**d**) this function in the two-dimensional section z = 0. (**e**) The approximation to the first negative and (**f**) to the first positive ripples of *G*(**x**) by the weighted functions *Ω*(**x**; *μ, v*) according to Table 1. Two-dimensional section z = 0 is shown. (**g**) The radial component of the function *Ω*(**x**; *μ, v*) with different *μ* and *v* values indicated on the plot. (**h**) The radial component of the difference between the left and right sides in the decomposition (1) for different numbers *M* of the terms included.

Differently from several approaches tried previously, *e*.*g*., (*4,5,13*-*18*), our method, giving accurate and analytically calculated map values, is based on the observation that both kinds of the principal map distortions, a resolution cut-off and a harmonic disorder, can be expressed by a convolution of an atomic density with a respective function. In the first case, it is the spherically symmetric interference function *G*(**x**) expressed through a dimensionless variable **x** = **r***D*^−1^ with the parameter *D* equal to the resolution cut-off in the Fourier space. In the second case, it is a normalized Gaussian function *g*(**x**; *v*) parameter *v* of which describes the scale of disorder and is proportional to the value known in structural biology as an atomic displacement parameter, *B* (see Supplementary materials).

The key to the analytic calculation of model maps is that the interference function can be decomposed into a linear combination

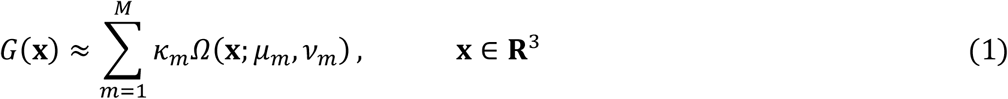

of terms

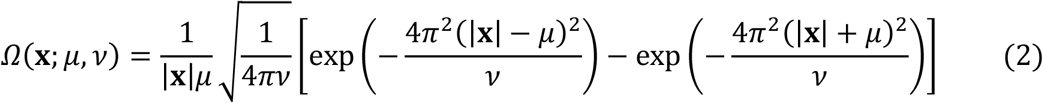

Each term describes the uniform distribution at the surface of the sphere of the radius *μ* convoluted with the Gaussian function *g*(**x**; *v*) and represents a solitary spherical wave, a ripple. The parameter *μ*_*m*_ in (1) corresponds to the radius of the *m*-th ripple, *v*_*m*_ reflects its width and *k*_*m*_ its height (Fig. 2e-2g), positive or negative. The number *M* of terms in (1) depends on the required cut-off distance and the accuracy of the decomposition (Fig. 2h). For *μ* = 0, function (2) coincides with the Gaussian function *g*(**x**; *v*).

Function (2) has been designed so that, being convoluted with a Gaussian function *g*(**x**; *v*_0_), it does not change its form but only substitutes its parameter *v* by *v* + *v*_0_. The last property means that an image of any ‘Gaussian atom’ 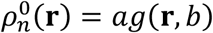 at any resolution *D*_*n*_ and with any displacement factor *B*_*n*_ is

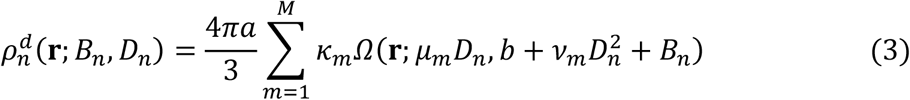

(Supplement materials). The values *μ*_*m*_, *v*_*m*_, *k*_*m*_ are parameters of the shell-decomposition (1), universally calculated for the required accuracy (*e*.*g*., Table 1). The parameter *B*_*n*_ manifests the disorder of the *n*-th atom while *D*_*n*_ describes the features of the experimental map in its local environment. These values can be refined together with the atomic position **r**_*n*_, depending on the amount of the experimental data available.

**Table 1.** Coefficients of the shell-decomposition (1) of the function *G*(**x**) obtained for |**x**| ≤ 10, *M* = 21 (Fig. 2h). Only three digits after comma are shown. The maximal error of the approximation is about 0.02% of the maximum of *G*(**x**) equal to 1. The central peak of *G*(**x**) is represented by the sum of the Gaussian function *k*_1_*Ω*(**x**; 0, *v*_1_) = *k*_1_*g*(**x**; *v*_1_) and the correcting function *k*_2_*Ω*(**x**; *μ*_2_, *v*_2_). Each Fourier ripple is represented by one *k*_*m*_*Ω*(**x**; *μ*_*m*_, *v*_*m*_) function alternating the sign of *k*_*m*_.

| $m$ | $\mu$ | $\nu$ | $\kappa$ | $m$ | $\mu$ | $\nu$ | $\kappa$ | $m$ | $\mu$ | $\nu$ | $\kappa$ |
| --- | --- | --- | --- | --- | --- | --- | --- | --- | --- | --- | --- |
| 1 | 0.000 | 10.131 | 0.693 | 8 | 3.492 | 2.795 | 0.485 | 15 | 6.989 | 1.670 | -0.368 |
| 2 | 0.339 | 3.216 | 0.026 | 9 | 3.971 | 2.882 | -0.476 | 16 | 7.490 | 1.509 | 0.356 |
| 3 | 0.873 | 4.819 | -0.797 | 10 | 4.471 | 2.022 | 0.401 | 17 | 7.991 | 1.369 | -0.334 |
| 4 | 1.439 | 3.622 | 0.595 | 11 | 4.995 | 1.620 | -0.371 | 18 | 8.493 | 1.248 | 0.326 |
| 5 | 1.979 | 3.616 | -0.599 | 12 | 5.504 | 2.317 | 0.416 | 19 | 8.995 | 1.146 | -0.332 |
| 6 | 2.462 | 4.143 | 0.623 | 13 | 5.980 | 2.062 | -0.407 | 20 | 9.494 | 1.060 | 0.333 |
| 7 | 2.953 | 3.047 | -0.534 | 14 | 6.490 | 1.849 | 0.392 | 21 | 9.978 | 0.811 | -0.290 |

In different experimental methods, the contribution 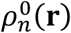 of an immobile atom to the exact density (or equivalently its scattering function) is usually approximated by a weighted sum of a few Gaussian functions with the coefficients *a*^(*k*)^, *b*^(*k*)^ tabulated for each type of atoms (*10,11*). Then decomposition (3) is naturally generalized for all types of atoms by introducing an outer sum over these Gaussian functions and can be used at any conventional resolution. At a subatomic resolution, a single-Gaussian model helps to describe a loss of spherical symmetry in 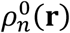 due to density deformation (*19*).

When the variation of the resolution may be neglected either over the whole molecule or locally, over a region of interest, the shell-decomposition can be used even more efficiently. First, one calculates numerically the image 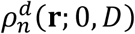 of an immobile atom of every required type at the given resolution *D*. Then the shell-decomposition (3) is built directly for each of these few oscillating images obtained with the exact scattering functions; this removes a need in any approximation to them. For each atom of the model, the only further adjustment required is increasing the values of the respective parameters *v*_*m*_ by the atomic *B*_*n*_; this gives their images very accurately (Table 2). The number of terms to calculate *ρ*^*d*^(**r**) is reduced in comparison with the general scheme accelerating the calculations and improving iterative refinement procedures further.

**Table 2.** Maximal error, up to the distance value *r*_*max*_, in the images of a carbon atom at the resolution of 2 Å calculated for different *B* values using the *Ω*-decomposition (3). The coefficients of the decomposition were calculated using the image of an immobile carbon atom with *B* = 0 up to the distance value *r*_*max*_ = 10.5 Å (Table S1). Second column shows, for comparison, the value of the image in the atomic center. The number in parentheses shows the number of terms sufficient to get such accuracy.

| $B, \text{Å}^2$ | $\rho^d(0)$ | Maximal error for $r \leq r_{max}$ ($M$ terms used) | | | |
| --- | --- | --- | --- | --- | --- |
|  |  | 2 Å (5) | 3 Å (6) | 4 Å (7) | 8 Å (12) |
| 0 | 1.97 | - | - | - | $1.9 \cdot 10^{-4}$ |
| 10 | 1.42 | $2.9 \cdot 10^{-6}$ | $5.3 \cdot 10^{-6}$ | $8.3 \cdot 10^{-6}$ | $8.6 \cdot 10^{-6}$ |
| 20 | 1.05 | $1.1 \cdot 10^{-6}$ | $4.0 \cdot 10^{-6}$ | $4.7 \cdot 10^{-6}$ | $4.8 \cdot 10^{-6}$ |
| 30 | 0.80 | $6.4 \cdot 10^{-6}$ | $2.9 \cdot 10^{-6}$ | $4.1 \cdot 10^{-6}$ | $4.3 \cdot 10^{-6}$ |

While such simplified version of the shell-decomposition seems to be useful at intermediate stages of model refinement, we expect that the full version with the refinement of *D*_*n*_ parameters would be important at the final stages. It will result in more accurate values of *B*_*n*_ parameters (*20*) separating their influence from that of the resolution cut-off (*12*), the value adjustable on fly. This in turn could allow, for example, extracting macromolecular dynamics from such data (*21*). Calculating such *Ω*-maps with all adjustable parameters, including resolution, will help to get model maps for the whole object; these maps, as accurate as possible, are required for their visual and numeric comparison with the experimental one. They are also important for model building and fixing eventual residual errors.

We illustrate an efficiency of our approach using a test protein model and calculated synthetic data allowing to shows the level of the method’s accuracy (Supplementary materials). We calculated an exact ‘diffraction map’ of a resolution of 2 Å for comparison and a series of the *Ω*-maps (3) with all *D*_*n*_ equal to 2 Å and with *M* that varied from one (the Gaussian peak only) to six. The map modeled ignoring the ripples, *M* = 1, was inaccurate in a number of regions while including a few first ripples by increasing *M* up to five made the differences with the exact map negligibly small (Fig. 1c). This proves a need in modeling the Fourier ripples which are ignored in a number of previous approaches.

To illustrate the possibility to reproduce a map of a prescribed inhomogeneous resolution, we assigned the resolution *D*_*n*_ varying from 2 Å in the model center to 5 Å at its periphery and calculated a respective model map. Indeed, the *Ω*-map was totally the same as the 2 Å – resolution Fourier map in the model center and coincided with the exact 5 Å – resolution Fourier map at the periphery. In between, where the local resolution was intermediate, the *Ω*-map was different from both control Fourier maps (see Supplementary materials for details).

The developed method has a number of features crucial for an efficient model refinement. Such refinement becomes the bottle neck to select hypothetical protein models suggested by structure prediction methods and to build other model components using MX or cryo-EM data. First, the method does not require any Fourier transform while can reproduce atomic images very accurately (Tables 1, 2; Fig. 1). Second, this method gives an analytic expression for the map values and for their derivatives with respect to all atomic parameters. The functions 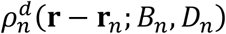 are given in the closed form and depend on the resolution and the displacement parameter analytically. Third, the most important and original, our method does not only make it trivial to calculate, in a single run, a map with the local resolution that varies from one region to another (Fig. 1b) but also to adjust this resolution on fly, to refine it according to the experimental map in the environment of a given atom and to deposit it. While such information is hardly available nowadays from cryo-EM databases such as EMDB (*22*) or EMPIAR (*23*), this will be trivial using the suggested approach.

Even when the decomposition (3) is illustrated by examples in cryo-EM and MX, it may be applied to oscillating spherically symmetric functions in other fields of science, in particular to calculate electrostatic potential in small- and macromolecular charge density studies (*24, 25*).

## Acknowledgments

We thank L. Urzhumtseva for her help with computing and A. Ben Shem and B. Klaholz for stimulating discussions of the results and the manuscript. AU thanks the French Infrastructure for Integrated Structural Biology (FRISBI) ANR-10-INSB-05-01 and Instruct, which is part of the European Strategy Forum on Research Infrastructures (ESFRI) and is supported by national member subscriptions.

## Funding

This research received no external funding.

## Author contributions

AU and VYL conceived of the theory, designed the numeric experiments and wrote the manuscript. AU developed the programs and performed numeric experiments.

## Competing interests

Authors declare that they have no competing interests.

## Data and materials availability

The code to calculate the shell-decomposition of atomic images, the atomic model used for the tests and the respective maps are available by request from the authors.

## Supplementary Materials

## Materials and methods

### S1. Maps and resolution

The term *resolution* is defined in structural biology quite ambiguously (*29*). When an experimental map is presented as a Fourier series, the resolution cut-off of this series is defined as the shortest period of the Fourier harmonics included into the calculation, and is traditionally considered as the *map resolution* (the number of Fourier harmonics eventually missed is supposed to be negligibly small).

To mimic the limited resolution of the experimental map, one may start by calculating the exact density distribution *ρ*(**r**) from the model followed by obtaining its Fourier coefficients. In crystallography, they are known as structure factors. The required map *ρ*^*d*^(**r**; *D*) is then calculated by the inverse Fourier transform with the set of structure factors limited by the resolution cutoff *D*. This procedure requires two Fourier transforms, does not provides one with simple analytic expressions for the derivatives of the score function, and does not allow getting a map of an inhomogeneous resolution, as those in cryo-EM. Instead, one may calculate such map as a sum of atomic contributions 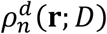 of the respective resolution. Several known approximations 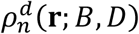 to atomic images (*4, 5,13-18*) either model only the central peak, or use image interpolations by *B* at a given resolution *D*, or use a step-function approximation to the atomic scattering function and the integrals of this approximation at a chosen resolution.

Variation of the local resolution, for example in cryo-EM, is usually illustrated by colored maps. It is inconvenient to conserve such information making it hardly depositing to and recovering from structural data bases.

### S2. Modeling principal map distortions

#### S2.1. Image of a function at a limited resolution

The image *ρ*^*d*^(**r**; *D*) of a function *ρ*(**r**) at the resolution *D* is defined as the integral over a sphere of the radius *D*^−1^

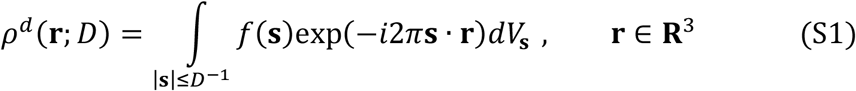

Here

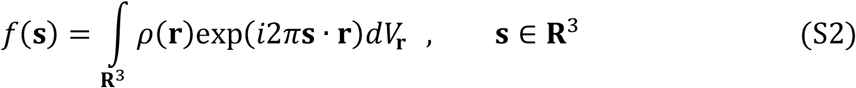

is the Fourier transform of the function *ρ*(**r**) calculated over the whole three-dimensional space. When *ρ*(**r**) is an atomic density, *f*(s) is known as the atomic scattering factor. We can rewrite (S1) as the integral over the whole space

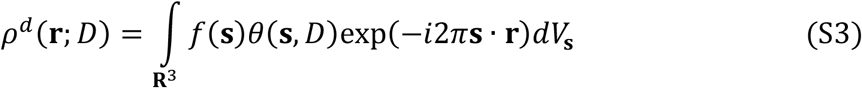

with a step-function

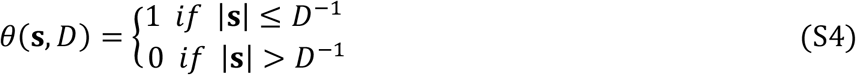

The integral (S3) can be presented as the convolution

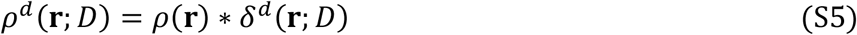

of *ρ*(**r**) with the spherically symmetric function

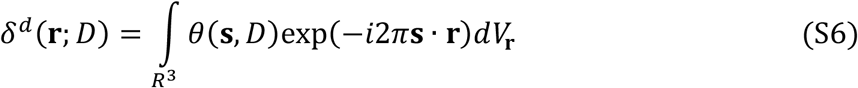

The last integral may be interpreted as the image of the delta-function *δ*(**r**) (or of an ‘immobile point atom’) at the resolution *D*.

The function (S6) can be calculated in the close form

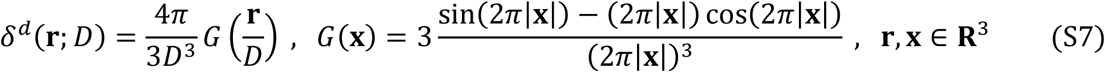

We express it through a spherically symmetric function *G*(**x**) of a dimensionless parameter

**x** = *D*^−1^**r**, the vector **r** rescaled by the resolution *D* (Fig. 2c,d). The function *G*(**x**) has a large peak in the origin surrounded by a number of spherically symmetric positive and negative ripples in space.

#### S2.2. Harmonic atomic disorder

A harmonic disorder of atomic positions in time and in space, individual for each atom, is usually modeled by a convolution of the respective contributions with a Gaussian function in three-dimensional space, in the simplest case the isotropic function:

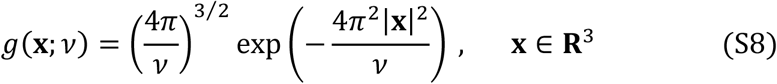

Such Gaussian model has a convenient computational feature. Its combination with another source of a harmonic disorder does not change its form but simply modifies the value of its parameter:

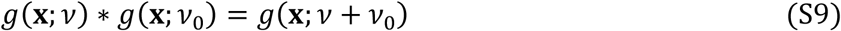

Since the convolutions are commutative, we can combine two principal sources of image distortion in the most convenient order.

### S3. Shell-decomposition of oscillating functions in space

#### S3.1. Uniform distribution at a spherical surface

A density of a ‘point atom’ with a harmonic disorder, described by a convolution of the Gaussian function with the delta-function, results in the same Gaussian. When, instead of the delta-function, we convolute this Gaussian function with the uniform distribution at the spherical surface of the radius *μ*, we obtain

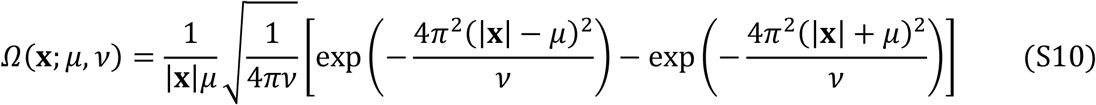

To get this function, we start from the uniform distribution in a thin spherical shell of the radius *μ* and width Δ, and then apply the convolution theorem and take the limit Δ→ 0. The Gaussian function, *g*(**x**; *v*), is a limit of *Ω*(**x**; *μ, v*) when *μ* → 0.

Function *Ω*(**x**; *μ, v*) has a number of properties similar to those for the Gaussian functions, in particular

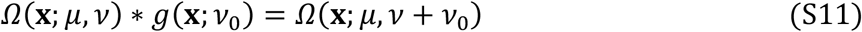

Another important feature is that the function conserves its form when rescaled from the dimensionless argument, **r** = *D***x**:

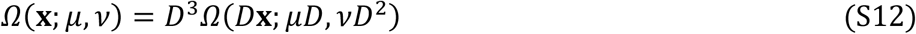

#### S3.2. Modeling three-dimensional ripples

Function *G*(**x**) does not have a property similar to (S11). To overcome this obstacle, we represent this function by a linear combination of functions (S10)

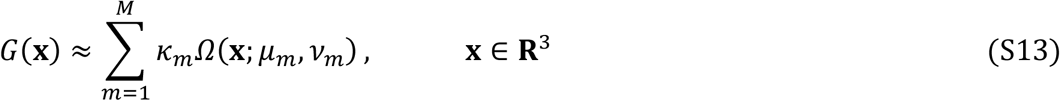

going beyond the approximation of its central peak by three-dimensional Gaussian functions (*13,14*) (terms with *μ*_*m*_ = 0). The optimal values of the parameters *μ*_*m*_, *v*_*m*_, *k*_*m*_ (Table 1) can be obtained minimizing the difference between two sides in (S13).

#### S3.3. Modeling atomic images

Combining (S7), (S12) and (S13) we obtain the image of an ‘immobile point atom’ at the resolution *D* as

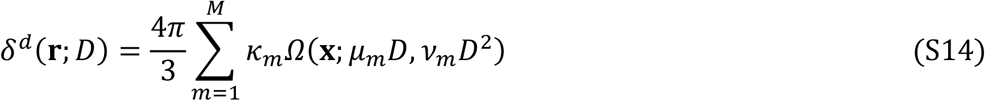

Similarly, we obtain an analytic expression for an image of a virtual ‘Gaussian atom’, or a ‘point atom with a harmonic isotropic disorder’ described by the atomic displacement parameter *B*_*n*_,

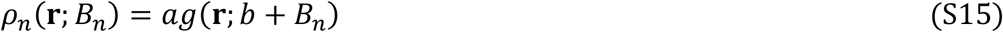

Since

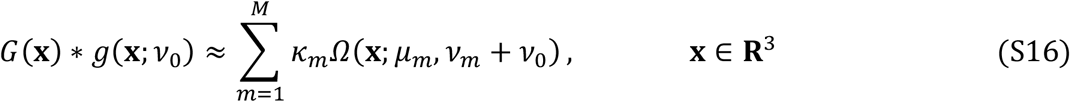

the image of a Gaussian atom (S15) in the position **r**_*n*_ is

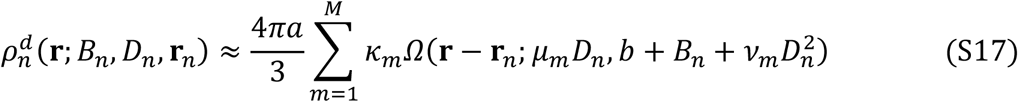

Except *B*_*n*_, *D*_*n*_ and the coordinates of the atomic center **r**_*n*_, all to be refined, all parameters in (S17) are external and known in advance. Analytical expression (S17) allows one to obtain simple analytic formulae for the derivatives of the map values with respect to all variable parameters. As a consequence, this gives the exact value of the gradient of the function that expresses the discrepancy between the experimental and model maps. High accuracy of this gradient assures that real-space refinement will results in the most accurate values of all model parameters including *B*_*n*_ and *D*_*n*_.

For actual, material atoms their potentials or densities are described by a sum of Gaussians with different (*a, b*) values (*10,11*). This introduces the additional outer sum in (S17) over these few Gaussians, all with the same atomic parameters and *μ, v, k* constants.

### S4. Models and data

For our tests, we took a medium-size atomic model from PDB (*1*) and calculated its electron density map at the resolution of 2 Å using the standard three-steps procedure with the double Fourier transform:

- calculation of the function *ρ*(**r**) as the sum of atomic densities *ρ*_*n*_(**r** − **r**_*n*_; *B*_*n*_);
- calculation of the structure factors by the Fourier transform of the function *ρ*(**r**);
- calculation of the map *ρ*^*d*^(**r**; *D*) by the inverse Fourier transform with the set of obtained structure factors limited by the resolution cutoff *D*.

The standard five-Gaussians approximation to the atomic scattering factors was used (*10*). This map was considered as the control one; Fig. 1c shows it at the 3σ cut-off. Then alternatively, we calculated a series of the *Ω*-maps (3) with different *M* values. The atomic contribution was calculated up to the distance of 5 Å from the atomic centers. Overall, the maps were similar to the control map. However, the map with *M* = 1, *i*.*e*., with no ripples included, was inaccurate in a number of regions.

The same model was used for the second test as described in the main text. Here both control maps, at the resolutions of 2 Å and 5 Å, were calculated using the standard three-step procedure described above.

---

**Table S1.** Coefficients of the shell-decomposition (3) of the image of a carbon atom at the resolution of 2 Å for |**r**| ≤ 10.5 Å, *M* = 12.

| $m$ | $\mu$ | $\nu$ | $\kappa$ | $m$ | $\mu$ | $\nu$ | $\kappa$ |
| --- | --- | --- | --- | --- | --- | --- | --- |
| 1 | 0.000000 | 41.609995 | 11.747566 | 7 | 5.965867 | 9.016016 | -5.200085 |
| 2 | 0.793310 | 9.990229 | 0.293693 | 8 | 6.980623 | 8.321347 | 4.992793 |
| 3 | 1.798971 | 18.459649 | -9.893183 | 9 | 7.982692 | 8.315746 | -4.924291 |
| 4 | 2.906666 | 14.085440 | 7.148594 | 10 | 8.974138 | 7.602500 | 4.714901 |
| 5 | 3.939076 | 12.155872 | -6.280142 | 11 | 9.866578 | 4.885647 | -2.064002 |
| 6 | 4.950601 | 10.384368 | 5.670299 | 12 | 10.103054 | 6.891677 | -2.449353 |

## Notes

### Competing Interest Statement

The authors have declared no competing interest.

